# Infralimbic activity during REM sleep facilitates fear extinction memory

**DOI:** 10.1101/2024.01.20.576390

**Authors:** Jiso Hong, Kyuhyun Choi, Marc V. Fuccillo, Shinjae Chung, Franz Weber

## Abstract

Rapid eye movement (REM) sleep is known to facilitate fear extinction and play a protective role against fearful memories. Consequently, disruption of REM sleep after a traumatic event may increase the risk for developing PTSD. However, the underlying mechanisms by which REM sleep promotes extinction of aversive memories remain largely unknown. The infralimbic cortex (IL) is a key brain structure for the consolidation of extinction memory. Using calcium imaging, we found in mice that most IL pyramidal neurons are intensively activated during REM sleep. Optogenetically suppressing the IL activity during REM sleep within a 4-hour window after auditory-cued fear conditioning impaired extinction memory consolidation. In contrast, REM-specific inhibition of the IL cortex after extinction learning did not affect the extinction memory. Whole-cell patch-clamp recordings demonstrated that inactivating IL neurons during REM sleep depresses their excitability. Together, our findings demonstrate that REM sleep after fear conditioning facilitates fear extinction by enhancing IL excitability, and highlight the importance of REM sleep in the aftermath of traumatic events for protecting against traumatic memories.

## INTRODUCTION

Persistent fear responses triggered by cues associated with a traumatic event are symptomatic of post-traumatic disorder (PTSD). PTSD patients show an impairment in learning to suppress fear responses to a previously fear-inducing stimulus that no longer predicts danger^1^, suggesting deficits in fear extinction^2^. Another core symptoms of PTSD are disturbances in rapid eye movement (REM) sleep with recurrent nightmares^3,4^. Pre-existing or persisting REM sleep disturbances after traumatic events significantly increase the risk for developing PTSD^5,6^, suggesting an involvement of REM sleep in fear processing^7,8^. In particular, growing evidence indicates an important role of REM sleep in extinction of conditioned fear^9,10^. Deprivation of REM sleep in humans and rodents impaired fear extinction in cued fear conditioning tasks^11–13^. Depleting REM sleep by ablating its braintem core circuitry in rodents similarly resulted in a weakened extinction memory^14^. However, the underlying mechanisms and the involved brain areas by which REM sleep facilitates fear extinction remain largely unknown.

The ventromedial prefrontal cortex (vmPFC) in humans and its rodent homolog, the infralimbic (IL) cortex^15^, is crucial for the consolidation of extinction memory, which suppresses fear responses to a conditioned stimulus^16^. The activity of the mPFC including the IL is strongly increased during REM sleep in mice^17^, and imaging studies in humans similarly revealed a strong activation of the vmPFC during REM sleep^18,19^. Imaging studies in PTSD patients identified structural abnormalities^20,21^ and hypoactivation of the vmPFC^22,23^ indicating that its dysfunction contributes to PTSD symptoms. Studies in rodents showed that electrolytic lesion^24^, pharmacological^25,26^ or optogenetic inhibition^27^ of the IL prior to or during extinction learning impairs extinction memory. In contrast, electrical stimulation^28,29^ or optogenetic activation of the IL enhanced extinction^27,30^, which is consistent with the facilitatory effect of transcranial direct current stimulation (tDCS) in humans^31^. Electrophysiological studies showed that fear conditioning depresses the excitability of IL neurons, while extinction reverses its excitability^32^. Pharmacologically or chemogenetically increasing IL excitability, in turn, facilitated extinction memory consolidation^33,34^, whereas its reduction resulted in impaired extinction^35,36^. Recent studies demonstrated that the REM sleep-specific activity of distinct neural populations plays an important role in the consolidation of spatial^37–39^ and social memories^40^, and deprivation of REM sleep can alter the intrinsic excitability of neurons^41,42^ and synaptic plasticity^43^. However, to what extent REM sleep affects the excitability of the IL and thereby possibly enhances extinction remains unknown.

Here, using calcium imaging, we found that the large majority of IL neurons are strongly activated during REM sleep. Suppression of IL activity specifically during REM sleep within 4 hours after conditioning reduced the excitability of IL neurons and weakened the recall of extinguished fear. These results indicate a crucial role of REM sleep within a specific time window in consolidating extinction memories.

## RESULTS

### IL pyramidal neurons are strongly activated during REM sleep

In our previous study, we showed that the bulk calcium activity of pyramidal neurons in the medial PFC (mPFC) including the IL is strongly increased during REM sleep^17^. However, the proportion of IL cortical neurons which were most active during REM sleep or other brain states remains unknown. To address this question, we performed cellular-resolution calcium imaging of pyramidal neurons in the IL using microendoscopy (**Fig. 1a,b**). Adeno-associated viruses (AAVs) encoding the fluorescence calcium indicator GCaMP6f (AAV-DIO-GCaMP6f) were injected together with AAV-CaMKII-Cre into the IL to monitor the calcium activity of pyramidal neurons (**Fig. 1a,b**). Imaging was performed through an implanted gradient refractive index (GRIN) lens connected to a miniaturized fluorescence microendoscope in freely moving mice (Fig. 1a, **Supplementary Fig. 1a,b**, and **Supplementary Video 1**). The imaged neurons were classified into three different subclasses depending on their average activity during each brain state (**Fig. 1c**). Strikingly, the majority of the IL pyramidal neurons (76.6%) were most active during REM sleep (REM-max, **Fig. 1d** and **Supplementary Fig. 1c**). Neurons with highest activity during wakefulness (Wake-max) comprised most of the remaining neurons; only one neuron was classified as NREM-max. (**Fig. 1d** and **Supplementary Fig. 1c**; see **Supplementary Table 1** for detailed statistical results). In sum, our calcium imaging experiments demonstrate that the large majority of IL pyramidal neurons are strongly activated during REM sleep.

**Figure 1.**
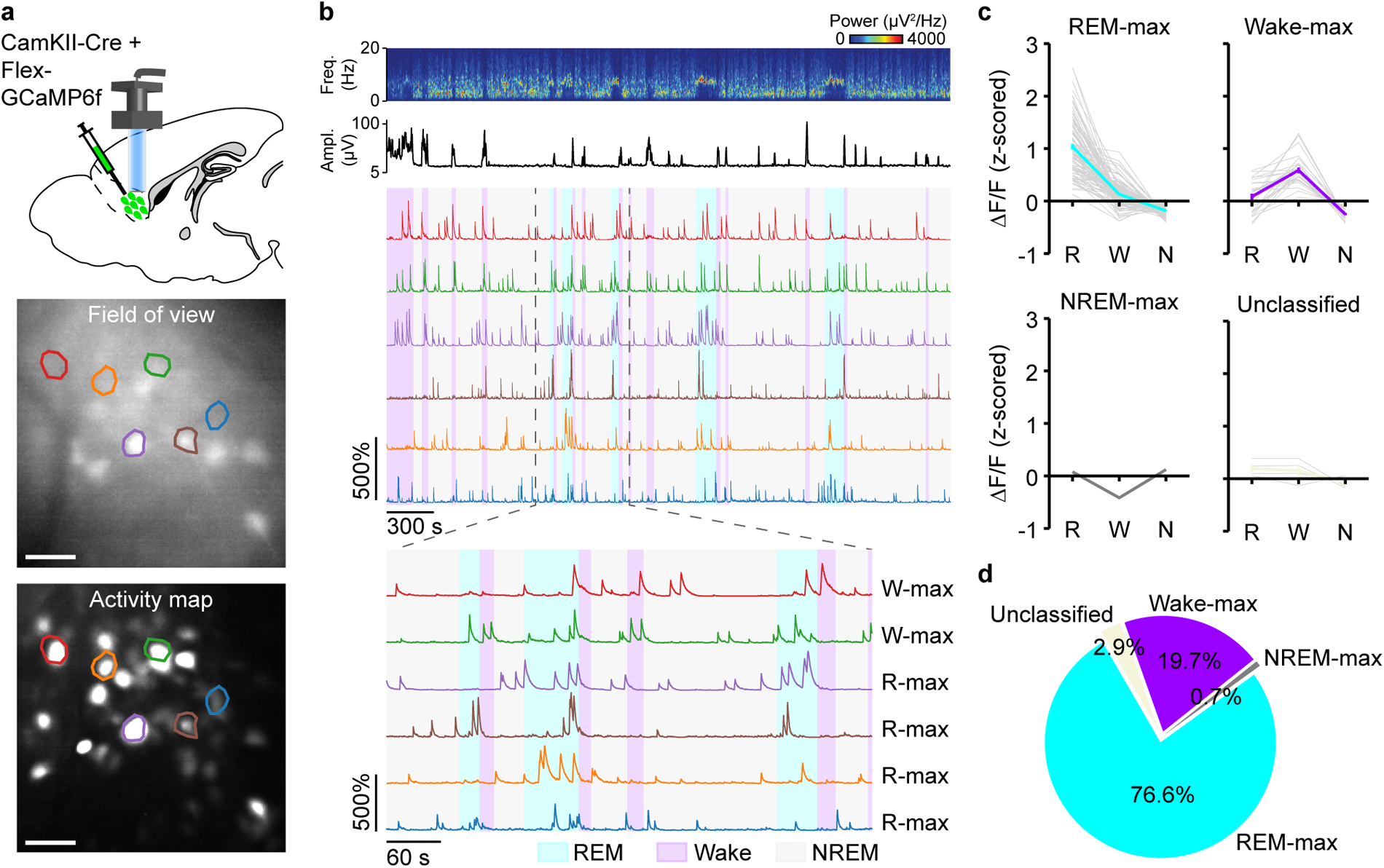
IL pyramidal neurons are strongly activated during REM sleep. (a) Top, experimental scheme of in vivo calcium imaging of IL pyramidal neurons. Bottom, field of view and activity map of an example imaging session. Colored polygons, example region of interest (ROIs). Scale bar, 50 µm. (b) Top, EEG spectrogram, EMG amplitude, and ΔF/F traces of the representative ROIs outlined in (a). Bottom, ΔF/F signals on an expanded timescale for a selected interval (indicated by dashed lines). (c) Average ΔF/F activity of different neuron subclasses for each brain state (R-max, n = 105; W-max, n = 27; N-max, n = 1; Unclassified, n = 4; n = 6 mice). Bold lines, mean across cells ± s.e.m.; gray lines, individual cells. (d) Proportion of different neuron subclasses.

### IL Inhibition during post-conditioning REM sleep impairs fear extinction memory

As the IL is known to be crucial for successful fear extinction^2,44^, we wondered whether its strong activation during REM sleep plays a necessary role in the consolidation of extinction memory. To test this, we silenced the activity of IL pyramidal neurons specifically during REM sleep after fear conditioning in a first set of experiments (**Fig. 2a,b**). On the conditioning day, mice were introduced to auditory-cued fear conditioning, comprising 6 pairings of a 20-s tone (conditioned stimulus, CS) co-terminating with a 1-s electric foot shock (unconditioned stimulus, US; **Fig. 2b**). The next day, mice underwent extinction learning with 48 presentations of the CS without foot shock. One day later during recall, the extinction memory was tested by presenting 3 times the CS in the extinction context. We measured the level of fear by quantifying the percentage of time spent freezing during each tone. Immediately after conditioning, we monitored sleep for 8 hours based on electroencephalogram (EEG) and electromyography (EMG) recordings and inhibited IL activity specifically during REM sleep using a closed-loop stimulation protocol (**Fig. 2b**). For optogenetic inhibition, the light-activated chloride channel iC++ was expressed in pyramidal neurons and a dual fiber-optic cannula was bilaterally implanted into the IL (**Fig. 2a** and **Supplementary Fig. 2a**). Control animals were instead injected with eYFP. As soon as the onset of a spontaneous REM sleep episode was detected based on real-time analysis of the EEG and EMG (**Methods**), laser stimulation was initiated and lasted until the end of the episode. To test whether the effects of IL inhibition on extinction were specific to REM sleep, we subjected another group of iC++ mice to closed-loop inhibition. But instead of administering the laser during REM sleep, we delayed it by one minute after the end of each REM episode and delivered it for the same duration. As expected, laser stimulation exclusively overlapped with REM sleep in both the iC++ and eYFP group (**Fig. 2c** top and **Supplementary Fig. 2b**). In contrast, in the iC++ delay group the laser coincided with either wakefulness or NREM sleep (**Fig. 2c**). Consistent with our previous study^17^, the EEG θ power during REM sleep bouts with laser stimulation was reduced in iC++ mice compared with REM sleep in baseline recordings without laser (**Fig. 2c** and **Supplementary Fig. 2c**). Moreover, laser stimulation caused a reduction in the frequency of phasic θ events, transient increases in the θ amplitude and frequency, which are characteristic of phasic REM sleep ^45,46^. Optogenetically suppressing IL activity during REM sleep did not induce significant changes in the amount, duration, or frequency of REM sleep (**Supplementary Fig. 2e**), nor did it affect the amount of wakefulness or NREM sleep (**Supplementary Fig. 2f**).

**Figure 2.**
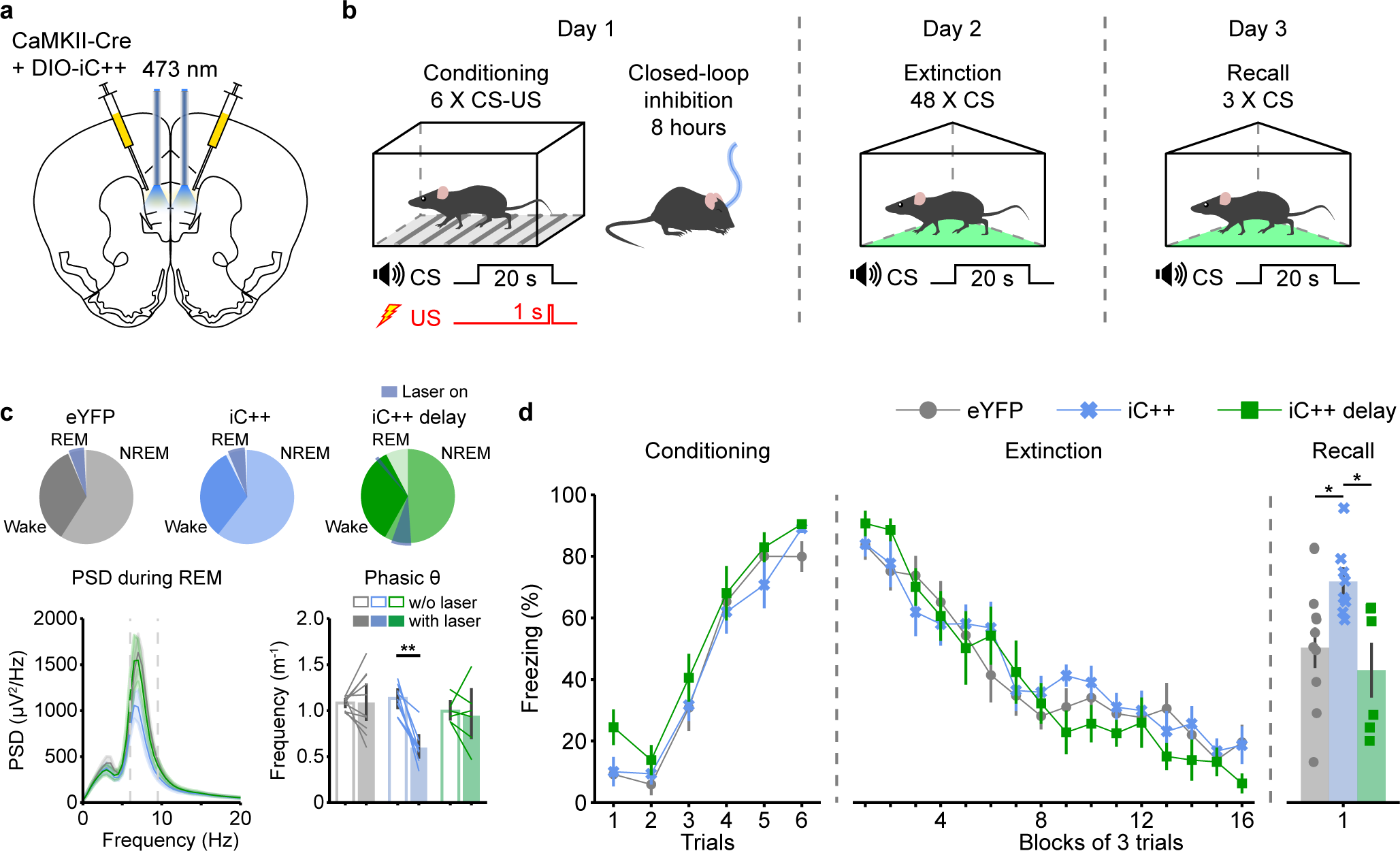
Closed-loop inhibition of IL pyramidal neurons during REM sleep after fear conditioning impairs extinction memory. (a) Schematic illustrating optogenetic inhibition of IL pyramidal neurons. (b) Experimental protocol for fear conditioning, extinction and recall with closed-loop inhibition after fear conditioning. Following fear conditioning, mice were transferred to their homecage and subjected for 8 hours to closed-loop inhibition of IL neurons specifically during REM sleep. (c) Top, amount of each brain state and overlap with laser application during sleep recordings after fear conditioning, represented as a pie chart. Bottom left, power spectral density (PSD) of the EEG during REM sleep. Dashed lines indicate the θ frequency range. Shadings, 95% CIs. Bottom right, frequency of phasic θ events during REM sleep episodes with closed-loop inhibition (filled bars) and episodes without inhibition (in baseline recordings without laser, empty bars). Error bars, 95% confidence intervals (CIs); lines, individual mice. Mixed ANOVA, laser, F(1, 21) = 11.3010, P = 0.0030; virus × laser interaction, F(2, 21) = 9.6629, P = 0.0011; pairwise tests, eYFP without vs with laser, T(9) = -0.0604, P = 1.0000; iC++ without vs with laser, T(7) = 6.4670, P = 0.0010; iC++ delay without vs with laser, T(5) = 0.4293, P = 1.0000; eYFP, n = 10; iC++, n = 8; iC++ delay, n = 6 mice. **P<0.001. (d) Freezing during auditory CS in fear conditioning, extinction, and recall. A block in extinction and recall comprises 3 consecutive tones. Error bars, ± s.e.m. Conditioning: mixed ANOVA, virus, F(2,21) = 2.1084, P = 0.1464; virus × trials interaction, F(10, 105) = 0.5501, P = 0.8506; extinction: mixed ANOVA, virus, F(2,21) = 0.1935, P = 0.8256; virus × blocks interaction, F(30, 315) = 1.2496, P = 0.1781; recall: one-way ANOVA, F(2, 21) = 5.3869, P = 0.0129; eYFP vs iC++, T(15.0284) = -2.9044, P = 0.0109; eYFP vs iC++ delay, T(10.1088) = 0.6982, P = 0.5008; iC++ vs iC++ delay, T(7.3724) = 3.0606, P = 0.0172; eYFP, n = 10; iC++, n = 8; iC++ delay, n = 6 mice. *P<0.05.

All three groups of animals showed successful acquisition of fear conditioning on day 1, as reflected in the increase in freezing across tones (**Fig. 2d**). The REM sleep-specific inhibition of the IL did not affect the strength of the fear memory (quantified by the percentage of freezing during the first three tones during extinction; **Supplementary Fig. 2d**) on day 2, and had no effect on extinction learning (**Fig. 2d**). Yet, interestingly, during the recall of extinction on day 3, iC++ mice exhibited higher freezing, suggesting that IL inhibition during post-conditioning REM sleep impairs the consolidation of the extinction memory (**Fig. 2d**). Given that the general sleep architecture in the iC++ group did not differ from that in the eYFP or iC++ delay groups (**Supplementary Fig. 2e,f**), the weakened extinction memory is likely the direct result of silencing IL activity during post-conditioning REM sleep and not the indirect consequence of changes in sleep as a result of laser stimulation. Altogether, our findings demonstrate that the IL activity during REM sleep after conditioning facilitates fear extinction memory.

### REM sleep’s facilitatory effect on fear extinction is time-dependent

To test whether the facilitatory effect of the IL activity on extinction is restricted to REM sleep within a specific time window, we conducted REM-specific IL inhibition starting 4 hours after fear conditioning ended in iC++ and eYFP control mice (**Fig. 3a,b** and **Supplementary Fig. 3a,b**). IL inactivation reduced the frequency of phasic θ events and the θ power during REM sleep (**Fig. 3c** and **Supplementary Fig. 3c**), but did otherwise not affect the sleep architecture (**Supplementary Fig. 3e,f**). Similar to IL inhibition performed directly after fear conditioning, REM-specific inhibition starting 4 hours later did not affect the fear memory (**Supplementary Fig. 3d**) or extinction learning (**Fig. 3c**). However, in contrast to IL inhibition after conditioning, starting laser stimulation 4 hours later did not result in significant differences in freezing between the eYFP and iC++ groups during the recall of fear extinction (**Fig. 3c**). These findings suggest that REM sleep occurring within 4 hours after fear conditioning plays a critical role in facilitating consolidation of the extinction memory, whereas modulation of REM sleep-specific IL activity outside this time window does not affect extinction.

**Figure 3.**
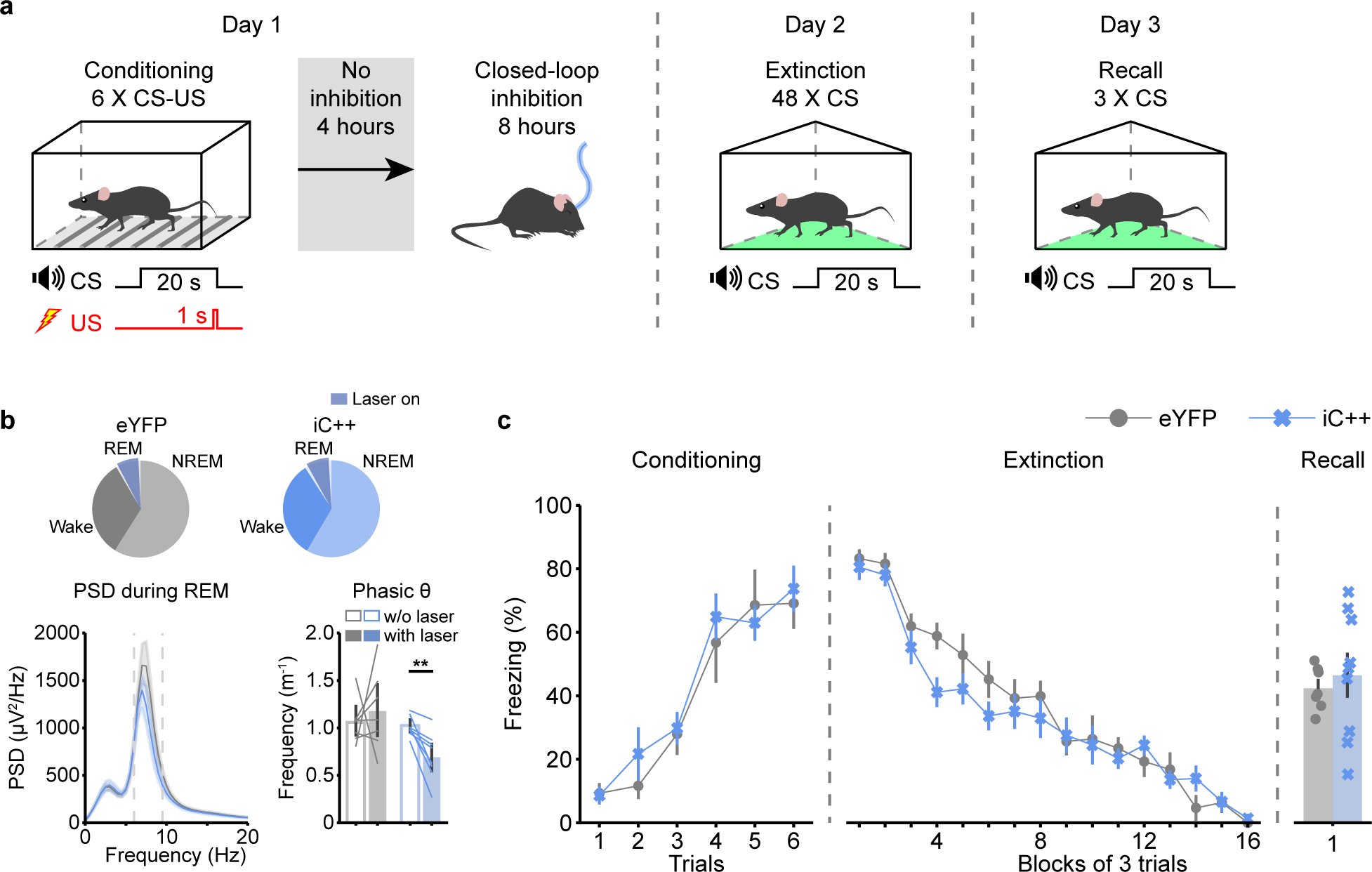
Closed-loop inhibition of IL neurons 4 hours after fear conditioning does not impair the extinction memory. (a) Experimental protocol for fear conditioning, extinction, and recall with closed-loop inhibition during REM sleep starting 4 hours after fear conditioning. (b) Top, amount of each brain state and overlap with laser application during sleep recordings starting 4 hours after fear conditioning. Bottom left, PSD of the EEG during REM sleep. Dashed lines indicate the θ frequency range. Shadings, 95% CIs. Bottom right, frequency of phasic θ events during REM sleep episodes with closed-loop inhibition (filled bars) and episodes without inhibition (in baseline recordings without laser, empty bars). Error bars, 95% CIs; lines, individual mice. Mixed ANOVA, laser, F(1, 14) = 1.7343, P = 0.2090; virus × laser interaction, F(1, 14) = 4.6903, P = 0.0481; pairwise tests, eYFP without vs with laser, T(6) = -0.5311, P = 0.6144; iC++ without vs with laser, T(8) = 4.9243, P = 0.0023; eYFP, n = 7; iC++, n = 9 mice. **P<0.01. (c) Freezing during auditory CS in fear conditioning, extinction, and recall. A block in extinction and recall comprises 3 consecutive tones. Error bars, ± s.e.m. Conditioning: mixed ANOVA, virus, F(1, 14) = 0.2306, P = 0.6385; virus × trials interaction, F(5, 70) = 0.5184, P = 0.7615; extinction: mixed ANOVA, virus, F(1, 14) = 1.9228, P = 0.1872; virus × blocks interaction, F(15, 210) = 1.3723, P = 0.1631; recall: t-test, T(10.2230) = 0.5716, P = 0.5799; eYFP, n = 7; iC++, n = 9 mice.

### Closed-loop inhibition of IL neurons during post-conditioning REM sleep reduces their excitability

Previous studies showed that the excitability of IL neurons during extinction learning correlates with the strength of the extinction memory^32,47^. We therefore examined the effects of closed-loop inhibition during post-conditioning REM sleep on the excitability of IL neurons. iC++ and eYFP animals were subjected to fear conditioning, followed by optogenetic closed-loop inhibition of IL pyramidal neurons during REM sleep (**Fig. 4a,b**). To specifically test the effects of REM sleep after conditioning on IL excitability, both cohorts did not receive extinction learning. Whole-cell patch-clamp recordings of IL neurons (**Supplementary Fig. 4a**) performed on day 3 showed that the resting membrane potential in iC++ cells was slightly depolarized compared with eYFP cells (**Supplementary Fig. 4b**), while the input resistance was not affected (**Supplementary Fig. 4c**). Interestingly, in response to depolarizing current steps with increasing amplitude, iC++-expressing cells fired fewer action potentials compared with eYFP-expressing cells (**Fig. 4c,d**), indicating that suppression of IL activity during post-conditioning REM sleep reduces the excitability of IL pyramidal neurons.

**Figure 4.**
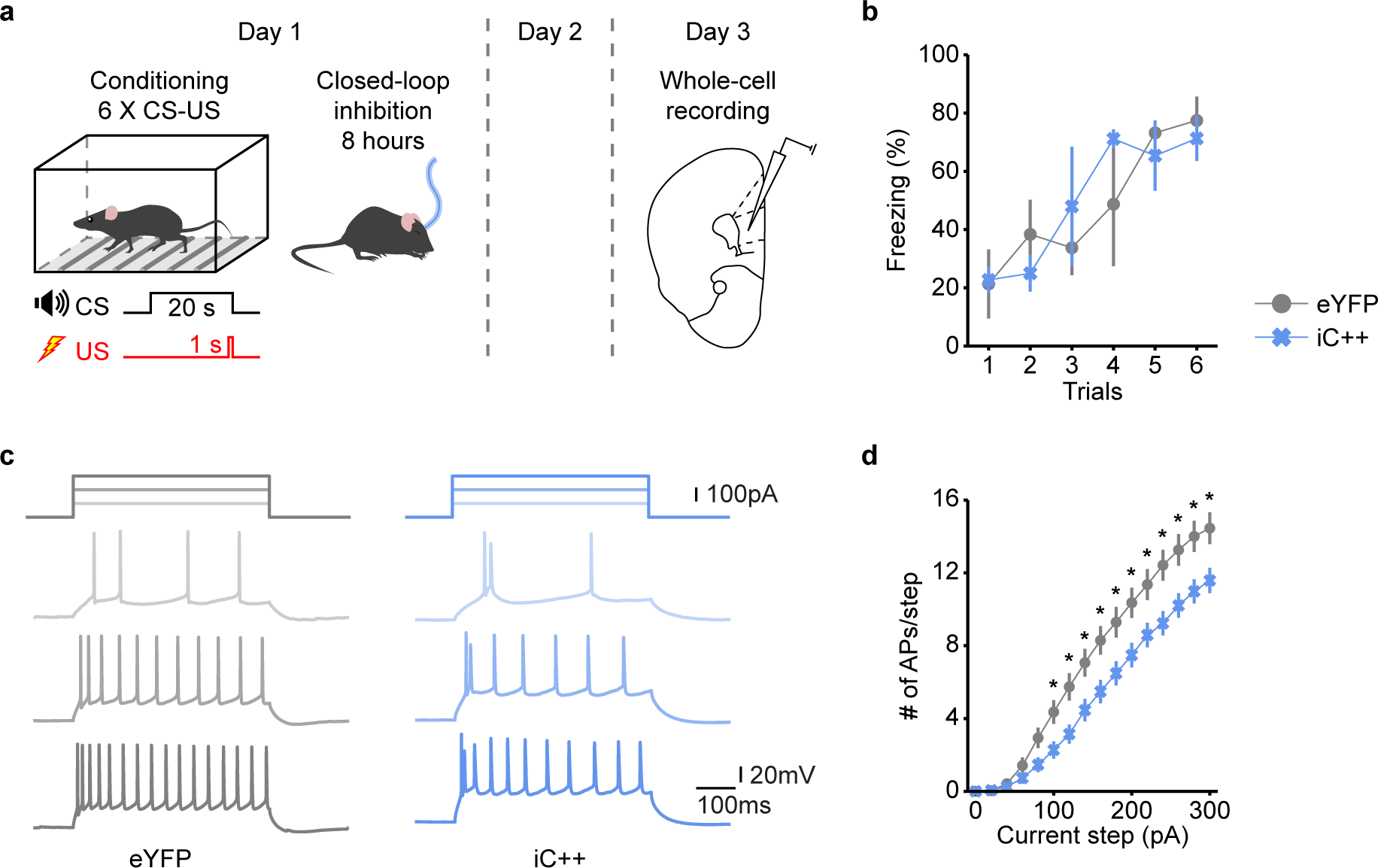
Closed-loop inhibition of IL neurons during REM sleep after fear conditioning reduces neural excitability. (a) Experimental protocol to assess the impact of REM sleep-dependent inhibition after fear conditioning on the excitability of IL neurons. (b) Freezing during each auditory CS in fear conditioning. Error bars, ± s.e.m. Mixed ANOVA, virus, F(1, 5) = 0.0511, P = 0.8301; virus × trials interaction, F(5, 25) = 0.8744, P = 0.5123; eYFP, n = 3; iC++, n = 4 mice. (c) Representative voltage traces of IL pyramidal neurons expressing eYFP (left) or iC++ (right) in response to depolarizing current pulses. (d) Number of action potentials evoked in response to current steps with different intensity. Error bars, ± s.e.m. Mixed ANOVA, virus, F(1, 77) = 10.5011, P = 0.0018; virus × current interaction, F(15, 1155) = 6.7495, P < 0.0001; eYFP, n = 31 cells from 3 mice; iC++, n = 48 cells from 4 mice. *P<0.05.

### Closed-loop inhibition during post-extinction REM sleep does not impair fear extinction

Lastly, we tested whether the IL activity during REM sleep after extinction learning is also involved in extinction memory consolidation. After conditioning and extinction acquisition, iC++ and eYFP control mice underwent closed-loop inhibition of IL neurons during REM sleep (**Fig. 5a,b** and **Supplementary Fig. 5a,b,d**). Inactivating the IL again reduced the θ power during REM sleep and the frequency of phasic θ events in iC++-mice (**Fig. 5b** and **Supplementary Fig. 5c**), but did not induce significant changes in sleep (**Supplementary Fig. 5e,f**). We found that freezing during extinction recall on day 3 did not differ between iC++ and eYFP mice (**Fig. 5c**), suggesting that IL activation during post-extinction REM sleep does not contribute to extinction memory consolidation.

**Figure 5.**
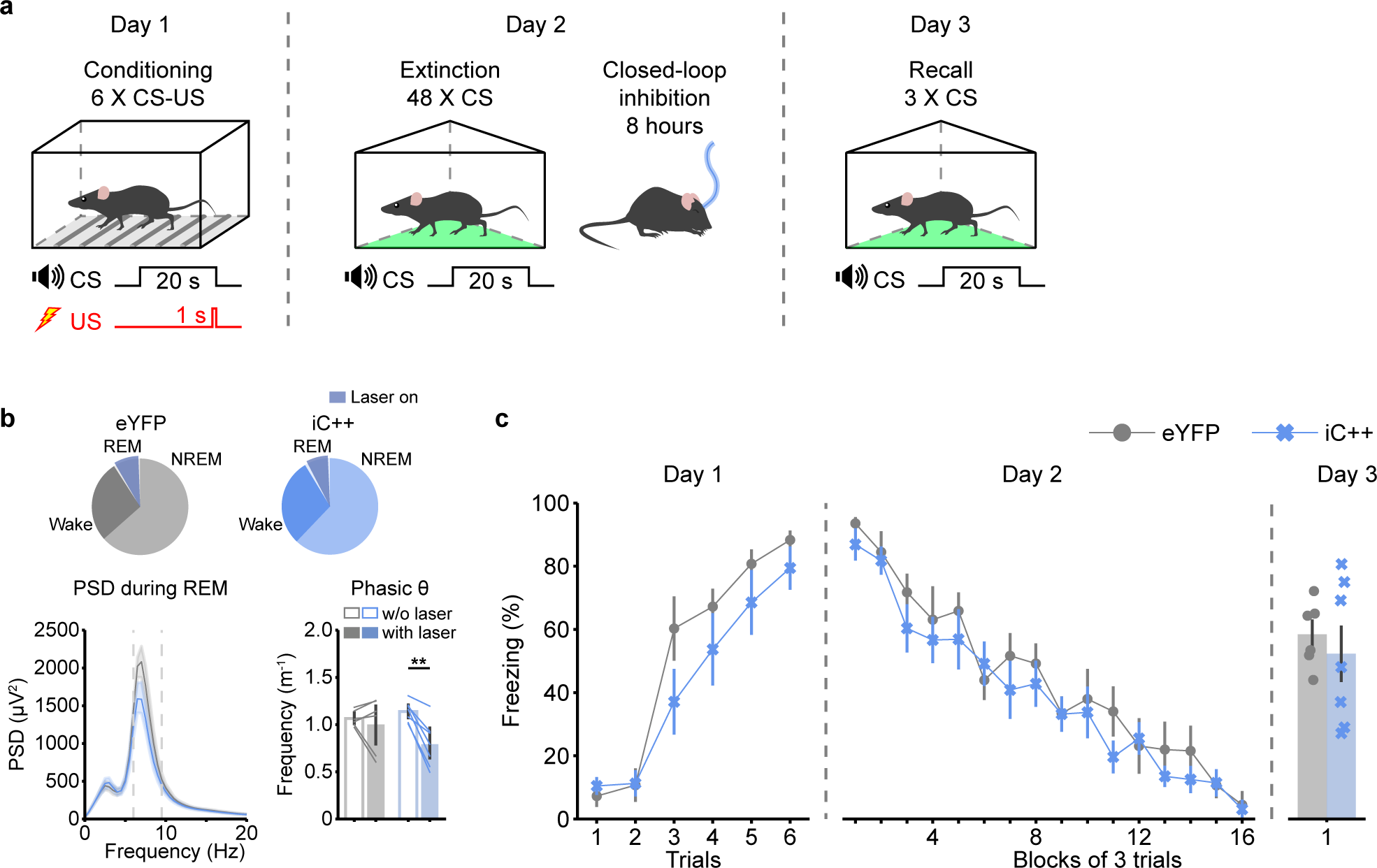
Closed-loop inhibition of IL neurons after fear extinction does not impair extinction memory. (a) Experimental protocol for fear conditioning, extinction, and recall with closed-loop inhibition of IL neurons after fear extinction. (b) Top, amount of each brain state and overlap with laser application during sleep recordings after fear extinction. Bottom left, PSD of the EEG during REM sleep after fear extinction. Dashed lines indicate the θ frequency range. Shadings, 95% CIs. Bottom right, frequency of phasic θ events during REM sleep episodes with closed-loop inhibition (filled bars) and episodes without inhibition (in baseline recordings without laser, empty bars). Error bars, 95% CIs; lines, individual mice. Mixed ANOVA, laser, F(1, 11) = 15.9511, P = 0.0021; virus × laser interaction, F(1, 11) = 6.9256, P = 0.0233; pairwise tests, eYFP without vs with laser, T(5) = 0.6775, P = 0.5282; iC++ without vs with laser, T(6) = 5.5515, P = 0.0029; eYFP, n = 6; iC++, n = 7 mice. **P<0.01. (c) Freezing during auditory CS in fear conditioning, extinction, and recall. A block during extinction and recall comprises 3 consecutive tones. Error bars, ± s.e.m. Conditioning: mixed ANOVA, virus, F(1, 11) = 1.6495, P = 0.2254; virus × trials interaction, F(5, 55) = 1.5676, P = 0.1846; extinction: mixed ANOVA, virus, F(1, 11) = 0.7654, P = 0.4003; virus × blocks interaction, F(15, 165) = 0.6362, P = 0.8418; recall: t-test, T(8.7247) = -0.6479, P = 0.5337; eYFP, n = 6; iC++, n = 7 mice.

## DISCUSSION

We found that the majority of IL pyramidal neurons are maximally activated during REM sleep. Suppressing IL pyramidal neurons specifically during REM sleep within a 4-hour window after fear conditioning impaired extinction memory consolidation. Inhibiting IL neurons during post-conditioning REM sleep reduced their intrinsic excitability, suggesting that REM sleep strengthens extinction memory by potentiating the excitability of IL pyramidal neurons.

The ventral and dorsal part of the mPFC are known to play functionally diverse roles in fear extinction^16^. In contrast to the IL, a recent study showed that the somatic activity of pyramidal neurons in the dorsal PFC is minimal during REM sleep^48^. Releasing their somatic inhibition by silencing PV neurons during post-conditioning REM sleep led to stronger fear memory and enhanced resistance to extinction learning. Thus, the opposing activity of the dorsal and ventral parts of the PFC during REM sleep reflects and may contribute to their different roles in fear processing.

Fear conditioning causes a depression in the excitability of IL neurons^32,49^. Successful extinction of fear and consolidation of extinction memory in turn requires the recovery of the IL neuron excitability, a process mediated by NMDA receptor-dependent synaptic plasticity^25,50^. The reduced IL excitability and impaired extinction memory by closed-loop inhibition during post-conditioning REM sleep (but not during NREM sleep or wake) indicates that this excitability modulation occurs during REM sleep. Supporting this notion, selective deprivation of REM sleep reduces neuronal excitability by impeding NMDA receptor function^42,51,52^, and application of an NMDA receptor agonist reversed the negative effects of REM sleep deprivation on extinction^53^. The increased cholinergic tone in the mPFC during REM sleep through the release of acetylcholine by the basal forebrain^54^ may further potentiate synaptic plasticity via NMDA receptor-dependent pathways^55,56^. Interestingly, injection of muscarinic agonists before extinction increased IL excitability and enhanced extinction memory^34^. Our results further suggest that REM sleep reverses the depression in IL excitability within a 4-hour time window after fear conditioning. Consistent with this finding, a recent contextual fear conditioning study showed that pharmacologically inhibiting the IL directly after conditioning, but not six hours later, made the fear memory more persistent to extinction^57^.

In our studies, closed-loop inhibition of IL neurons during REM sleep reduced the theta power and frequency of phasic θ events^17^. Hippocampal θ oscillations during REM sleep are involved in memory consolidation^58,59^, and their disruption by inhibiting the medial septum during REM sleep impaired spatial memory consolidation^37^. Phasic θ events support inter-regional coordination between cortical and hippocampal areas^45,60^, and their suppression by IL neuron inactivation may thus contribute to the impaired extinction memory.

We found that REM-specific IL inhibition after fear conditioning, but not after extinction, weakened the extinction memory. This finding aligns with research in humans showing that sleep deprivation after fear conditioning results in poor extinction memory^61,62^. Moreover, a recent study observed a positive relationship between the quantity of REM sleep preceding extinction learning and the strength of extinction memory in humans^63^. Likewise, studies in rats reported worse extinction memory when performing REM deprivation after conditioning in both cued and contextual fear tasks^64^. However, other studies also found impaired extinction memory when performing total sleep or REM sleep deprivation after extinction learning^65,66^. Importantly, our closed-loop approach specifically suppressed the REM-specific activation within the IL, while REM sleep deprivation globally disrupts the activity in brain regions. Therefore, the effects of post-extinction REM deprivation on extinction memory likely involve other brain areas or may result from the indirect effects of REM deprivation on other brain states^67^. Similarly, the reported strengthening of fear memories by REM sleep after fear conditioning is likely due to its impact on other emotion-regulatory brain areas such as the amygdala^13,68^.

Impairment in extinction memory consolidation has been proposed as a core mechanism underlying the development of PTSD^1^. Our findings suggest a potential explanation for the link between disrupted sleep, specifically abnormalities in REM sleep, following a traumatic event, and the subsequent development of symptoms of PTSD and anxiety disorders^5,6^. Sleep disruption subsequent to trauma might prevent the strong activation of the vmPFC during REM sleep, required to restore the excitability of vmPFC neurons. Our study underscores the importance of good REM sleep after a traumatic event and may explain the tenacity of PTSD symptoms if pharmacological or behavioral intervention occurs too late.

## ACKNOWLEDGEMENTS

This work was supported by a NARSAD Young Investigator grant (27799) by the Brain & Behavior Research Foundation, and by a grant from the Margaret Q. Landenberger Foundation to FW. We thank Justin Baik for help with setting up the sleep recording system, Caroline Johansen, Stephanie Acquaye, and Camille Harrison for help with histology and sleep annotation.

## AUTHOR CONTRIBUTIONS

Conceptualization: JH, FW; Funding Acquisition: FW; Investigation: calcium imaging and behavioral experiments: JH; patch clamp recordings: KC; Project Administration: FW, SC, MVF; Data analysis: JH; Writing: JH, FW.

## DECLARATION OF INTERESTS

The authors declare no competing financial interests.

## METHODS

### Animals

All animal care and experimental procedures were approved by the Institutional Animal Care and Use Committee (IACUC) at the University of Pennsylvania and conducted in accordance with the National Institutes of Health Office of Laboratory Animal Welfare Policy. Adult male C57BL/6J mice aged 8 - 9 weeks old at the point of surgery were used for all experiments. Food and water was available *ad libitum* under a 12:12 hr light:dark cycle with light on from 7:00 to 19:00. The colony room was maintained at an ambient temperature of 20 - 23°C and humidity of 40 - 60%. Mice for auditory fear conditioning and closed-loop inhibition were group-housed except during sleep recordings. Animals for patch clamp recordings were single-housed after fear conditioning until the electrophysiological recordings were performed. Mice for microendoscope imaging were single-house after GRIN lens implantation.

### Surgical procedures

All surgeries were performed following the IACUC guidelines for rodent survival surgery. Mice were anesthetized with 1-4% (vol/vol) of isoflurane in oxygen after subcutaneous injection of meloxicam (5 mg per kg body weight) or meloxicam ER (2 mg per kg body weight) and positioned in a stereotaxic frame (David Kopf Instruments) with heating pad for maintaining the body temperature. The skin was incised to gain access to the skull after asepsis.

For EEG recordings, two screws were bilaterally fixed on the top of the parietal cortex, and stainless-steel wires (A-M systems) were attached to them. For EMG recordings, two stranded stainless-steel wires (A-M systems) were inserted into the neck muscles. The reference electrode was connected to a screw located on top of the left cerebellum. All electrodes were connected to a mini-connector prior to the surgery and secured to the skull with dental cement.

Virus injections were performed using Nanoject II (Drummond Scientific) with a glass micropipette inserted into the IL (anteroposterior (AP) +1.85 mm; mediolateral (ML) ± 0.35 mm; dorsoventral (DV) -2.35 mm). For calcium imaging of pyramidal neurons, AAV9-CaMKII0.4-Cre-SV40 (0.1 μl, 2.9 x 10^13^ genome copies (GC)/ml, University of Pennsylvania vector core) and AAV1-Syn-Flex-GCaMP6f-WPRE-SV40 (0.4 μl, 1.3 x 10^13^ GC/ml, Addgene) was injected into the IL. 14 to 21 days later, a GRIN lens (500 μm diameter, Inscopix) was implanted into the IL (DV -2.3 mm). For optogenetic inhibition, AAV9-CaMKII0.4-Cre-SV40 (0.1 μl, 2.9 x 10^13^ GC/ml) and AAV2-Ef1a-DIO-iC++-eYFP (0.4 μl, 5.9 x 10^12^ GC/ml, University of North Carolina vector core) was bilaterally injected into the IL. Implantation of a dual fiber-optic cannula (200 μm diameter, Doric lenses) followed virus injection. Mice with no virus expression, where virus expression was outside the target region, or where the optic fiber was misplaced were excluded from the dataset.

### Histology

Mice were deeply anesthetized and transcardially perfused with 0.1M PBS and 4% paraformaldehyde (vol/vol) and brains were removed. After overnight fixation in paraformaldehyde, brains were stored in 30% sucrose in PBS (wt/vol). For histology, 40 μm sections cut using a cryostat (Thermo Scientific HM525NX) were placed onto glass slides (Fisher Scientific) and mounted with Fluoromount-G (SouthernBiotech) and cover glass (Globe Scientific). Fluorescence images were taken using a fluorescence microscope (microscope, Leica DM68; camera, Leica DFC7000GT).

### Auditory fear conditioning and extinction

Fear conditioning, extinction learning, and recall were performed during the light cycle, 8:00 to 10:00. Auditory fear conditioning was performed in a square acrylic box with dimensions 18.5 x 20 x 30 cm (L×WxH), placed within a sound-attenuating chamber with lights on. The conditioning box was equipped with a stainless steel grid on the bottom connected to a current-regulated shocker (Coulborn Instrument) and a USB video camera (Actimetrics, 15-Hz frame rate) attached to the top of the box to record the animals’ behavior. Stimulus (auditory tone or electric shock) presentation and EEG/EMG signal recordings were controlled by the Synapse software (Tucker-Davis Technologies) and a TDT RZ5P amplifier (Tucker-Davis Technologies, samplicate 1.5 kHz). On day 1, mice were placed in the conditioning box and habituated for 3 min. After habituation, mice received six tones (CS, 3000 Hz, 80 dB, 20 s each) co-terminated with an electric shock (US, 0.5 mA, 1 s), separated by 2 min intervals. 60 s after the presentation of the CS-US pairings, mice were moved to their recording cages for sleep recordings. The conditioning box was cleaned with 70% EtOH between each session. For fear extinction on day 2 and recall on day 3, mice were placed in a different context, i.e., a triangular acrylic box (27 cm on each side) with white paper bedding, located within a dark chamber. The animal behavior was recorded with a USB infrared camera (ELP, 15 Hz frame rate) mounted above the box. After 3 min of habituation, mice received 48 tones (extinction) or 3 tones (recall) interspaced by 40 s intervals. 60 s after the final tone presentation, mice were transferred to their recording cages for sleep recordings and REM sleep-specific optogenetic inhibition. After each session, the extinction/recall box was cleaned with water, and the bedding in the box was exchanged.

### Polysomnographic recordings with REM sleep-dependent closed-loop inhibition

To test the role of REM sleep-dependent IL activation on extinction memory, we subjected different groups of mice to the following REM sleep-specific optogenetic inhibition protocols: (1) Closed-loop inhibition during REM sleep during 8-hour EEG/EMG recordings starting immediately after fear conditioning, (2) closed-loop inhibition starting one min after each REM sleep episode and lasting for the same duration as the preceding REM sleep episode (**Fig. 2**), (3) closed-loop inhibition during REM sleep during 8-hour recordings starting 4 hours after fear conditioning (**Fig. 3**), and (4) closed-loop inhibition during 8-hour recordings starting after fear extinction (**Fig. 4**).

Prior to sleep recordings, each mouse was habituated to its individual recording cage located within a sound-attenuating chamber for 2 days. Sleep recordings with optogenetic inhibition were performed during the light cycle after fear conditioning or extinction. Baseline recordings without laser stimulation were performed one day before fear conditioning. EEG and EMG signals were recorded via a flexible recording cable using an RHD2132 amplifier (Intan Technologies, sampling rate 1 kHz) connected to the RHD USB Interface Board (Intan Technologies), which was controlled by the RHD Recording Controller software (Intan Technologies, version 1.5.2). In addition, videos were recorded using a camera (Chameleon3, FLIR, 5-Hz frame rate) mounted above the mouse cage.

For closed-loop inhibition, a flexible patch cable connected with a blue laser (Laserglow) was attached to the implanted optic ferrule in addition to the EEG/EMG recording cable. The timing of laser pulses was controlled using TTL signals generated by a Raspberry Pi (https://github.com/justin0bk/socketrecv/), which was controlled by a custom-programmed user interface (https://github.com/justin0bk/sleepRecording_v9/). REM sleep was detected automatically based on real-time spectral analysis of EEG and EMG signals. For the REM sleep detection, thresholds for the δ power and EMG amplitude were calculated for each mouse based on baseline recordings as well as a hard and soft threshold for the θ/δ ratio. The onset of a REM sleep bout was defined as the time point when the δ power and EMG amplitude were lower than their respective thresholds and when the θ/δ ratio surpassed the hard threshold. Step-pulse laser stimulation (3.0 - 3.5 mW at fiber tip) lasted until the end of the REM sleep bout, defined as the time point when the θ/δ ratio dropped lower than the soft threshold or till the EMG amplitude surpassed its threshold.

### Electrophysiology

Prior to the recording, mice were subjected to auditory fear conditioning and closed-loop inhibition during REM sleep for 8 hours on day 1. Electrophysiological recordings of IL neurons were performed on day 3. Mice were deeply anesthetized and trans-cardially perfused with ice-cold aCSF containing (in mM): 124 NaCl, 2.5 KCl, 1.2 HaH_2_PO_4_, 24 NaHCO_3_, 5 HEPES, 13 Glucose, 1.3 MgSO_4_, 2.5 CaCl_2_. After perfusion, the brain was quickly removed, submerged and coronally sectioned on a vibratome (VT1200s, Leica) at 250 μm thickness in ice-cold aCSF. Slices were transferred to NMDG based recovery solution at 32°C of the following composition (in mM): 92 NMDG, 2.5 KCl, 1.2 NaH_2_PO_4_, 30 NaHCO_3_, 20 HEPES, 25 Glucose, 5 Sodium ascorbate, 2 Thiourea, 3 Sodium pyruvate, 10 MgSO_4_, 0.5 CaCl_2_. After 12-15 minutes recovery, slices were transferred to room temperature aCSF chamber (20-22°C) and left for at least 1 hour before recording. Following recovery, slices were placed in a recording chamber, fully submerged at a flow rate of 1.4∼1.6 mL/min, and maintained at 29-30°C in oxygenated (95% O2, 5% CO2) aCSF.

For current-clamp recordings, the recording pipette was filled with internal solution containing (in mM) 140 K-gluconate, 5KCl, 0.2 EGTA, 2 MgCl_2_, 10 HEPES, 4 MgATP, 0.3 NaGTP, 10 2Na-Phosphocreatine (pH adjusted to 7.3-7.4 using KOH). To evaluate the passive membrane properties of the cells, they were maintained at their resting membrane potential. A step-wise protocol was then applied for 500 ms, where each sweep involved an increase of +20 pA in current injection. This step-wise protocol continued until a maximum current injection of 300 pA was reached. Each sweep in the step-wise protocol was separated by a duration of 15 seconds.

Recordings were performed using a MultiClamp 700B (Molecular Devices) and Igor7 (WaveMetrics; recording artist addon, developed by Richard C Gerkin, github: https://github.com/rgerkin/recording-artist), filtered at 2.8 kHz and digitized at 10 kHz. Data were analyzed using Igor7.

### Microendoscopy imaging

For cellular resolution calcium imaging of pyramidal neurons in the IL cortex, 1 - 2 weeks after GRIN lens implantation, a base plate was fixed on the animal’s head on top of the lens using dental cement to attach the Miniscope V3.2 (Labmaker, Germany). Imaging sessions took place during the light cycle in the home cage placed within a sound attenuating chamber and lasted for 1.5 hours. For each animal, we performed one or two imaging sessions on separate days. The microendoscope was connected via a flexible cable to the Miniscope PCB and an additional flexible cable was attached to the mini-connector on the animal’s head for EEG/EMG recordings using an RHD2132 amplifier (1 kHz sampling rate). The mice were habituated to the recording system for at least 1 hour after attaching the Miniscope to the baseplate. The microendoscope camera was controlled using the Miniscope V3.2 data acquisition software (https://github.com/daharoni/Miniscope_DAQ_Software) and calcium imaging movies were acquired with a frame rate of 20 Hz.

### Imaging analysis and definition of different subclasses of IL neurons

To correct for lateral motion in the recorded calcium imaging videos, we created a spatially high-pass filtered image stack by subtracting from each image a spatially low-pass filtered version of itself. We then manually selected a high-contrast area within the mean projection of the high-pass filtered image stack as spatial reference. For each movie frame in the high-pass filtered stack, we performed a 2D cross-correlation to determine the shift in the x- and y-direction optimizing the overlap between the current movie frame and the reference. To select regions of interest (ROIs) for further analysis, we first computed an activity map, M_x,y_, highlighting pixels with strong variations in their intensity over time using

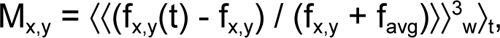

where f_x,y_(t) denotes the fluorescence at pixel (x,y) of movie frame t, f_x,y_ is the average of f_x,y_(t) over time, and f_avg_ refers to the average fluorescence across the whole recording. The notation ⟨…⟩_t_ depicts the temporal average, and ⟨…⟩_w_ is the output of a spatial 2x2 box filter. Cell body-shaped regions of interest (ROIs) on the activity map with high intensity were manually selected by encircling them with polygons using a custom-programmed graphical user interface. A raw fluorescence trace (F(t)) was extracted for each ROI as the average across pixels within that ROI for each frame. To correct for contamination of the fluorescence signal by out-of-focus neuropil^69^, we calculated for each ROI the neighboring neuropil signal, F_np_(t), within a bordering 10 μm broad ring with ∼5 μm distance from the perimeter of each ROI, excluding other ROIs, and subtracted F_np_(t), scaled by a correction factor c, from the raw ROI signal, F(t): F_subt_(t) = F(t) − c F_np_(t). The correction factor c was estimated for each recording session by calculating the ratio between the mean pixel intensity of a manually selected blood vessel, F_blood_vessel,_ and a nearby region lacking an ROI signal, F_near_blood_vessel_, each subtracted by the mean pixel intensity of an off-lens region, F_off-lens_, i.e., c = (F_blood_vessel_ − F_off-lens_) / (F_near_blood_vessel_ − F_off-lens_). The baseline of each neuropil subtracted fluorescence signal, B(t), was estimated by calculating the linear regression fit to the values of F_subt_(t) for periods of low fluorescence activity, defined as values within the 20th percentile of each recording session^70^. Using the baseline, we calculated the relative change in fluorescence as a function of time using ΔF/F(t) = (F_subt_(t) - B(t)) / B(t). Finally, we extracted for each ROI a denoised fluorescence trace using the OASIS algorithm with an AR(1) model^71^. To identify the same ROIs within two imaging sessions, we first determined for each recording session a map with all ROIs and then rotated and translated these maps with respect to each other such that the overlap between the ROIs was maximized. ROIs from two imaging sessions with on overlap of at least 50% were interpreted as the same neuron. Calcium imaging recordings with weak fluorescence signals or no clearly detectable cell-body shaped ROIs were excluded from the data set.

For each ROI, we tested whether its activity was significantly modulated by the brain state using one-way ANOVA (with ΔF/F values as dependent variable and brain state as factor). Using the Tukey’s HSD post-hoc test, we determined for each ROI that was significantly modulated by brain state (P < 0.05) during which state the ΔF/F activity was highest.

### Statistics and reproducibility

No statistical methods were used to predetermine sample sizes but our sample sizes were similar to those reported in previous publications^37,38^. The location of fiber implants and virus expression were histologically verified and consistent across all animals in an experimental group. Representative histology images thus reflect the findings for all animals belonging to the same group. Statistical analyses were performed using the Python module pingouin (https://pingouin-stats.org/). All statistical tests were two-sided and a P-value < 0.05 was considered as significant. Data were compared using t-tests or ANOVA followed by multiple comparisons tests (pairwise t-tests with Holm–Bonferroni correction for mixed ANOVA and Tukey’s post hoc test or Holm–Bonferroni correction for one-way ANOVA). Using the Shapiro–Wilk test, we verified that the data were normally distributed. For mixed ANOVA, Mauchly’s test was applied to check the sphericity of the data. In case sphericity was violated, P-values were corrected using the Greenhouse–Geisser correction. Statistical test results are included in the corresponding figure legends, Supplementary Information, and **Supplementary Table 1**.

## Notes

### Competing Interest Statement

The authors have declared no competing interest.

