## Supplementary Information for "Infralimbic activity during REM sleep facilitates fear extinction memory"

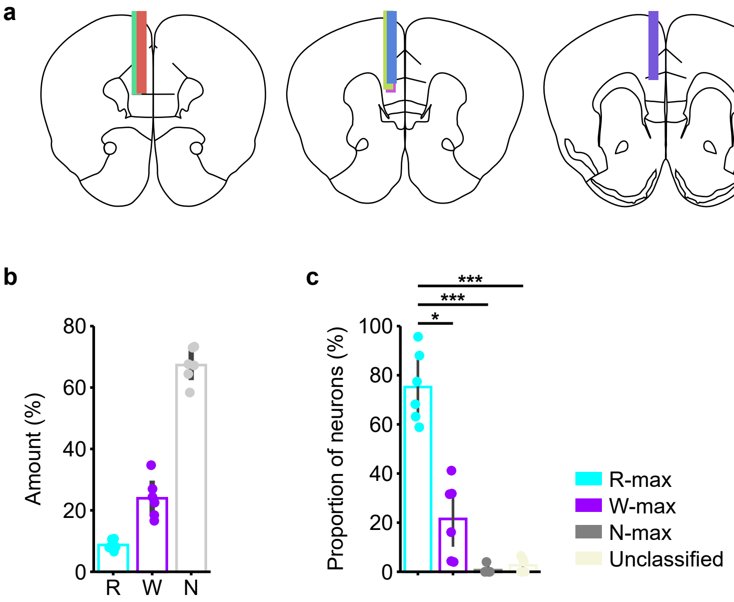


**Supplementary Fig. 1. Placement of GRIN lenses in IL, sleep architecture, and proportion of neuron subclasses.**

**(a)** Coronal sections depicting the location of GRIN lenses used to image pyramidal neurons in the infralimbic cortex (IL). n = 6 mice.

**(b)** Average amount of each brain state during calcium imaging experiments; R - REM, W - Wake, N - NREM. Error bars, 95% confidence intervals (CIs); dots, individual mice. n = 6 mice.

**(c)** Proportion of neuron subclasses (R-max, W-max, or N-max) across mice. Error bars, 95% CIs; dots, individual mice. One-way repeated measured ANOVA, $F(3, 15) = 46.1541$, $P = 0.0007$; t-tests with Holm-Bonferroni correction, R-max vs W-max, $T(5) = 4.3944$, $P = 0.0282$; R-max vs N-max, $T(5) = 13.1734$, $P = 0.0003$; R-max vs unclassified, $T(5) = 12.0445$, $P = 0.0003$; n = 6 mice. *$P<0.05$, ***$P<0.001$. See **Supplementary Table 1** for detailed statistical results.

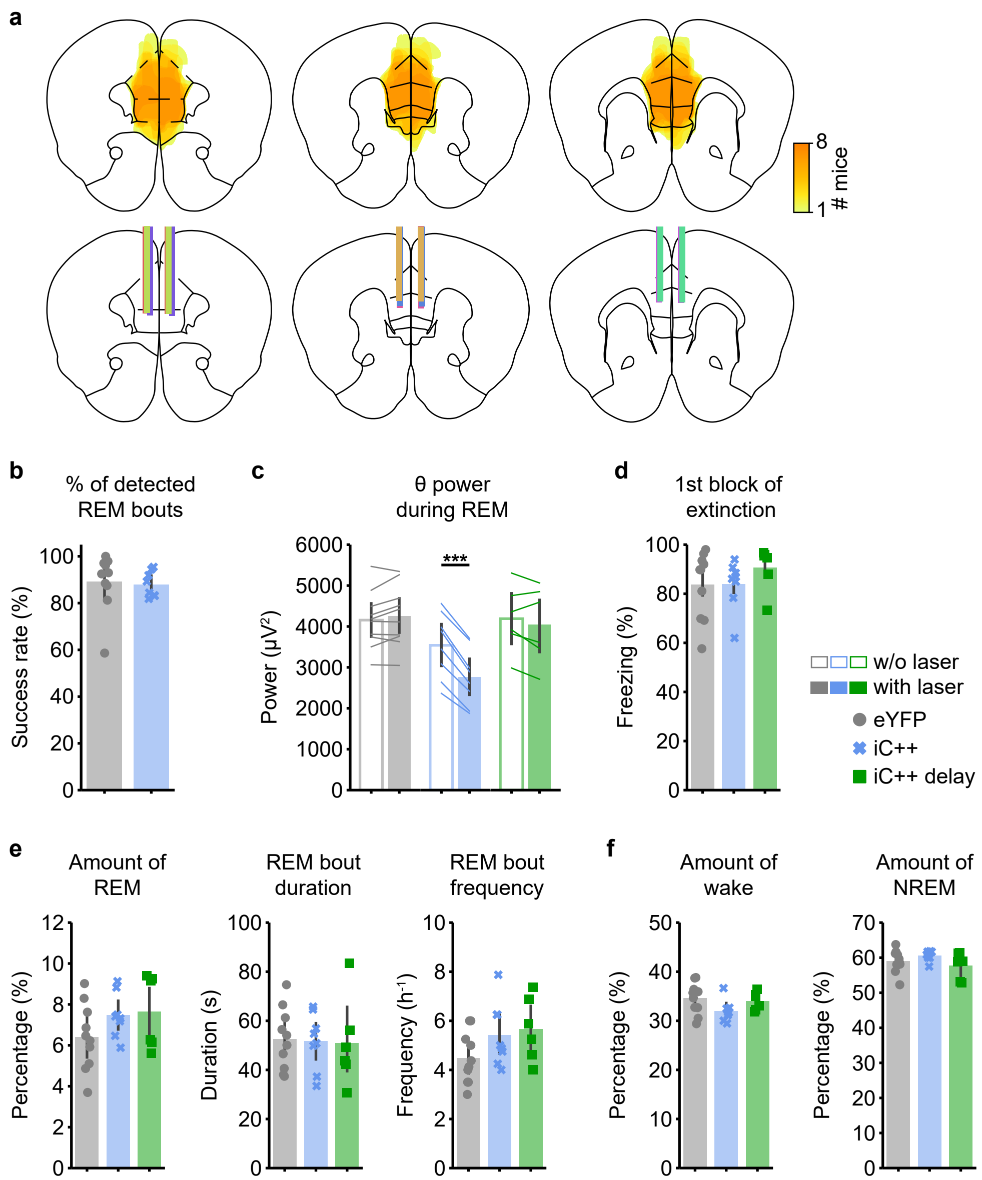


**Supplementary Fig. 2. Expression of iC++-eYFP and sleep in cohorts subjected to closed-loop inhibition directly after fear conditioning.**

**(a)** Top, heatmaps showing expression of iC++-eYFP in IL (iC++ group). Bottom, location of optic fiber tracts in IL.

**(b)** Percentage of REM bouts detected by closed-loop REM detection system during 8-hour recordings directly after fear conditioning. Error bars, 95% CIs; dots, individual mice. T-test, $T(13.0750) = 0.2957$, $P = 0.7721$; eYFP, $n = 10$; iC++, $n = 8$ mice.

**(c)** EEG power in θ range during REM sleep episodes with closed-loop inhibition (filled bars) and episodes without inhibition (in baseline recordings without laser, empty bars). Error bars, 95% CIs; lines, individual mice. Mixed ANOVA, laser, $F(1, 21) = 38.2300$, $P = 0.0000$; virus × laser interaction, $F(2, 21) = 43.3627$, $P = 0.0000$; pairwise tests, eYFP without vs with laser, $T(9) = -1.7565$, $P = 0.2258$; iC++ without vs with laser, $T(7) = 12.8536$, $P = 0.0000$; iC++ delay without vs with laser, $T(5) = 1.2993$, $P = 0.2505$; eYFP, $n = 10$; iC++, $n = 8$; iC++ delay, $n = 6$ mice. ***$P<0.001$.

**(d)** Mean freezing during the first 3 auditory CS presentations in fear extinction. Error bars, ± s.e.m. One-way ANOVA, $F(2, 21) = 0.7979$, $P = 0.4635$; eYFP, $n = 10$; iC++, $n = 8$; iC++ delay, $n = 6$ mice.

**(e)** Amount, average bout duration, and frequency of REM sleep during 8-hour interval after fear conditioning. Error bars, 95% CIs; dots, individual mice. Amount of REM sleep, one-way ANOVA, $F(2, 21) = 1.6614$, $P = 0.2139$; duration, one-way ANOVA, $F(2, 21) = 0.0287$, $P = 0.9717$; frequency, one-way ANOVA, $F(2, 21) = 2.2917$, $P = 0.1258$; eYFP, $n = 10$; iC++, $n = 8$; iC++ delay, $n = 6$ mice. **(f)** Amount of wake and NREM sleep during 8-hour interval after fear conditioning. Error bars, 95% CIs; dots, individual mice. Amount of wake, one-way ANOVA, $F(2, 21) = 2.2853$, $P = 0.1265$; amount of NREM, one-way ANOVA, $F(2, 21) = 1.5737$, $P = 0.2308$; eYFP, $n = 10$; iC++, $n = 8$; iC++ delay, $n = 6$ mice.

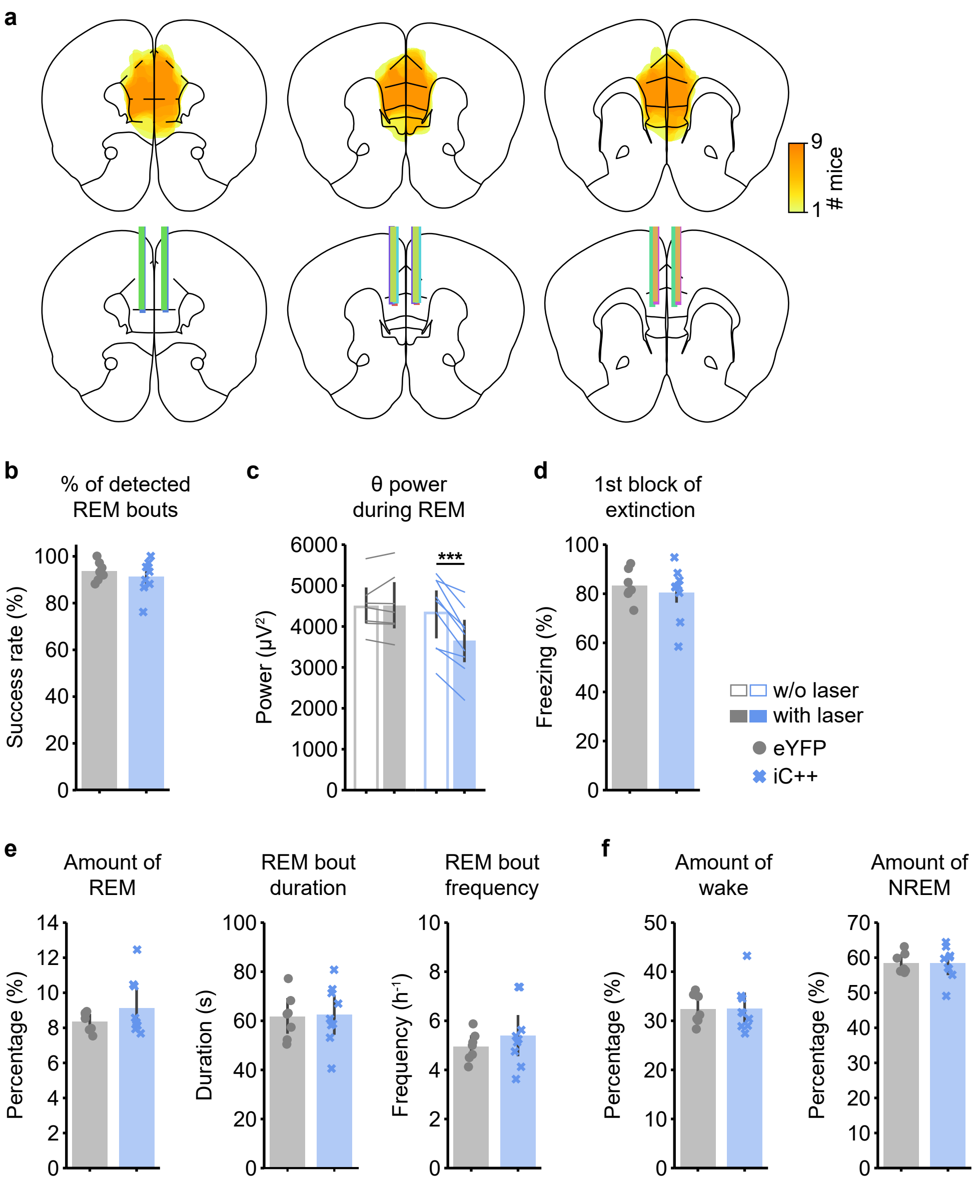


**Supplementary Fig. 3. Expression of iC++-eYFP and sleep in cohorts subjected to closed-loop inhibition starting 4 hours after fear conditioning.**

**(a)** Top, heatmaps showing expression of iC++-eYFP in IL. Bottom, location of optic fiber tracts in IL.

**(b)** Percentage of REM bouts detected by closed-loop system during 8-hour recordings starting 4 hours after fear conditioning ended. Error bars, 95% CIs; dots, individual mice. T-test, T(13.1352) = 0.8023, P = 0.4367; eYFP, n = 7; iC++, n = 9 mice.

**(c)** EEG θ power during REM sleep episodes with closed-loop inhibition (filled bars) and episodes without inhibition (in baseline recordings without laser, empty bars). Error bars, 95% CIs; lines, individual mice. Mixed ANOVA, laser, F(1, 14) = 28.4022, P = 0.0001; virus × laser interaction, F(1, 14) = 26.4236, P = 0.0002; pairwise tests, eYFP without vs with laser, T(6) = -0.4571, P = 0.6637; iC++ without vs with laser, T(8) = 6.3465, P = 0.0004; eYFP, n = 7; iC++, n = 9 mice. ***P<0.001.

**(d)** Mean percentage of time spent freezing during the first 3 CS presentations in fear extinction. Error bars, ± s.e.m. T-test, T(13.2956) = -0.6296, P = 0.5396; eYFP, n = 7; iC++, n = 9 mice.

**(e)** Amount, average bout duration, and frequency of REM sleep during 8-hour interval starting 4 hours after fear conditioning. Error bars, 95% CIs; dots, individual mice. Amount, t-test, T(10.2601) = -1.3319, P = 0.2117; duration, t-test, T(13.9950) = -0.1818, P = 0.8584; frequency, t-test, T(11.7418) = -0.9482,

P = 0.3621; eYFP, n = 7; iC++, n = 9 mice.

**(f)** Amount of wake and NREM sleep during 8-hour interval starting 4 hours after fear conditioning. Error bars, 95% CIs; dots, individual mice. Amount of wake, t-test, T(13.6572) = -0.0330, P = 0.9741; amount of NREM, t-test, T(13.5592) = -0.0071, P = 0.9944; eYFP, n = 7; iC++, n = 9 mice.

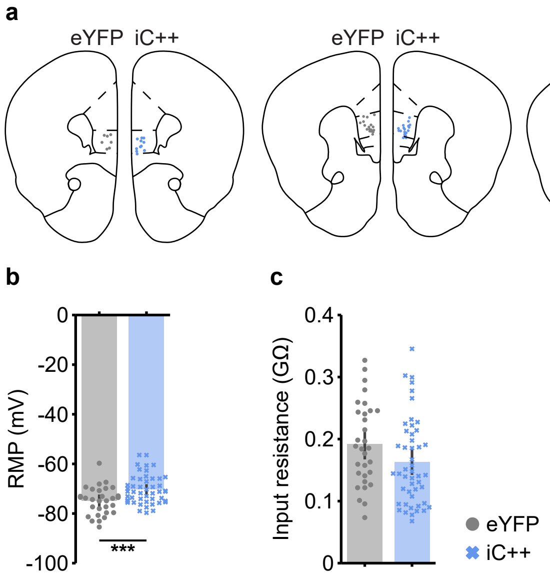


**Supplementary Fig. 4. Location, membrane potential, and input resistance of IL neurons.**
**(a)** Location of IL cells recorded in eYFP (left) and iC++ mice (right).
**(b)** Resting membrane potential (RMP) of recorded cells. Error bars, 95% CIs. T-test, T(66.5855) = -3.4713, P = 0.0009; eYFP, n = 31 cells from 3 mice; iC++, n = 48 cells from 4 mice. ***P<0.001.
**(c)** Input resistance of recorded cells. Error bars, 95% CIs. T-test, T(66.1903) = 1.9038, P = 0.0613; eYFP, n = 31 cells from 3 mice; iC++, n = 48 cells from 4 mice.

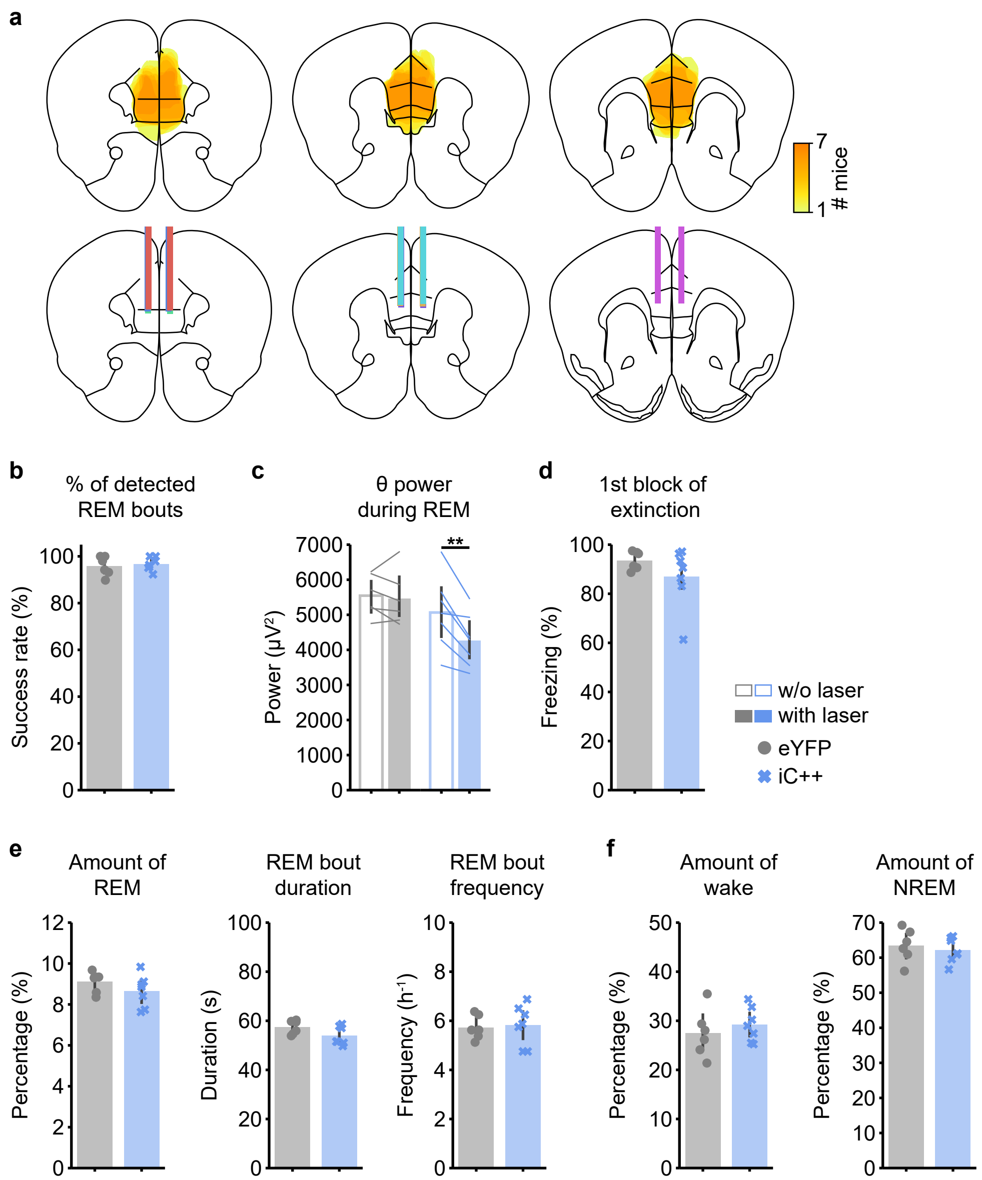


**Supplementary Fig. 5. Expression of iC++-eYFP and sleep in cohorts subjected to closed-loop inhibition directly after fear extinction.**

**(a)** Top, heatmaps showing expression of iC++-eYFP in IL. Bottom, location of optic fiber tracts in IL.

**(b)** Percentage of REM bouts detected by closed-loop system during 8-hour recordings directly after fear extinction. Error bars, 95% CIs; dots, individual mice. T-test, T(8.5704) = -0.4334, P = 0.6754; eYFP, n = 6; iC++, n = 7 mice.

**(c)** EEG θ power during REM sleep episodes with closed-loop inhibition (filled bars) and episodes without inhibition (in baseline recordings without laser, empty bars). Error bars, 95% CIs; lines, individual mice. Mixed ANOVA, laser, F(1, 11) = 15.2425, P = 0.0025; virus × laser interaction, F(1, 11) = 8.6874, P = 0.0133; pairwise tests, eYFP without vs with laser, T(5) = 0.5657, P = 0.5960; iC++ without vs with laser, T(6) = 4.4261, P = 0.0089; eYFP, n = 6; iC++, n = 7 mice. **P<0.01.

**(d)** Mean freezing during the first 3 CS presentations in fear extinction. Error bars, ± s.e.m. T-test, T(7.2712) = 1.3525, P = 0.2168; eYFP, n = 6; iC++, n = 7 mice.

**(e)** Amount, average bout duration, and frequency of REM sleep during 8-hour interval after fear extinction. Error bars, 95% CIs; dots, individual mice. Amount, t-test, T(10.3947) = 1.2337, P = 0.2445; duration, t-test, T(10.8636) = 1.8677, P = 0.0890; frequency, t-test, T(9.9509) = -0.2496, P = 0.8080;

eYFP, n = 6; iC++, n = 7 mice.

**(f)** Amount of wake and NREM sleep during 8-hour interval after fear extinction. Error bars, 95% CIs; dots, individual mice. Amount of wake, t-test, $T(8.9609) = -0.7544$, $P = 0.4700$; amount of NREM, t-test, $T(9.2192) = 0.5797$, $P = 0.5760$; eYFP, n = 6; iC++, n = 7 mice.

**Supplementary Table 1. Summary of statistical results for Figures and Supplementary Figures**

Each column in the table (Supplementary Table 1.xlsx) lists the

- Figure/Supplementary Figure reference ('Figure'),
- statistical test for group analysis ('Group analysis'),
- statistic and degrees of freedom (DOFs) for group analysis ('Statistic'),
- P-value for group analysis ('P-value'),
- effect size for group analysis ('Effect size'),
- comparisons ('Comparisons'),
- statistical test for pairwise tests ('Pairwise tests'),
- statistic and DOFs for pairwise tests ('Statistic'),
- P-values for pairwise tests ('P-value'),
- effect size for pairwise tests ('Effect size'),
- test type ('Tail'),
- sample size ('Sample size'),
- subjects ('Subjects').

**Supplementary Video 1. *In vivo* calcium imaging of IL pyramidal neurons.**
Calcium activity of IL pyramidal neurons recorded using a microendoscope (left) along with EEG spectrogram, EMG amplitude, and $\Delta F/F$ calcium traces. Several cells (colored ROIs) are drawn on top of the calcium video. The calcium traces (right) are represented using the same color code. The video is shown at a 30x speedup.
